# Single Cell Chemical Proteomics (SCCP) Interrogates the Timing and Heterogeneity of Cancer Cell Commitment to Death

**DOI:** 10.1101/2021.04.21.440805

**Authors:** Ákos Végvári, Jimmy E Rodriguez, Roman A Zubarev

## Abstract

Chemical proteomics studies the effects of drugs upon cellular proteome. Due to the complexity and diversity of tumors, the response of cancer cells to drugs is also heterogeneous, and thus proteome analysis at single cell level is needed. Here we demonstrate that single cell proteomics techniques have become quantitative enough to tackle the drug effects on the target proteins, enabling single cell chemical proteomics (SCCP). Using SCCP, we studied here the time-resolved response of individual adenocarcinoma A549 cells to anticancer drugs methotrexate, camptothecin and tomudex, revealing the early emergence of cellular subpopulations committed and uncommitted to death. As a novel and useful approach to exploring the heterogeneous response to drugs of cancer cells, SCCP may prove to be a “killer application” for single cell proteomics.

## 1. Introduction

Chemical proteomics studies the effects of drugs on cellular proteomes with the purpose of deciphering the targets and mechanisms of action (MOAs) of these molecules. ^1–3^ When sensitive cells are treated with toxic compounds for extended period of time, mechanistic target proteins become significantly regulated, and their profiling provides the first hint on the compound’s targets and MOA.^4,5^ This approach has been employed in functional identification of target by expression proteomics (FITExP)^6^, which laid ground for the online chemical proteomics ProTargetMiner tool.^1^ In FITExP, cells are treated at a LC_50_ concentration for 48 h, by which time half of the cells die. The dying cells detach from the substrate (for adherent cell types) and are found floating on the flask surface. In the remaining (surviving) cells, the drug target’s expression level is significantly and specifically regulated up or down, which serves a basis for drug target identification in FITExP. As an example, when cancer cells undergo treatment with methotrexate (MTX), the target protein dihydrofolate reductase (DHFR) becomes highly upregulated before the cells undergo programmed cell death.^7–9^ Interestingly, while the proteomes of the dying and surviving cells are very different (lending support to the notion that cell death is the ultimate case of cell differentiation), the drug target behaves in a similar manner in both types of cells. ^2^

The adherent cells usually start losing their attachment to the surface after 24 h of treatment at LC_50_ concentration, but the decision to survive or become dying must be made by the cell well before that.^1,10,11^ The intricate details of this decision-making process are of great scientific interest, as they possibly hold keys to the drug resistance mechanisms.^1,6^ These decision making processes can only be studied at the single cell level,^12^ while all so far reported chemical proteomics studies relied on bulk cell analysis.^13^ Cellular heterogeneity is currently analyzed routinely by single cell transcriptomics (SCT),^14^ with mRNA levels assumed to be proportional to the protein expression levels. However, at any given moment the concentration of both mRNA and proteins reflects the balance between their corresponding expression and degradation, and while mRNA transcription and protein expression are linked together rather well, the degradation processes for mRNA and proteins are completely decoupled. As a result, in the biological processes driven mostly by protein expression, mRNA levels provide excellent proxy for protein concentrations, but this correlation seems to break down already at steady states of the cell.^15^ In cell death processes mediated by protein degradation (*e*.*g*., via caspase proteases), a correlation between mRNA and protein levels cannot be presumed. Therefore, cell heterogeneity in death-related processes can best be studied with single cell proteomics (SCP).

Compared to the rather well developed SCT approaches, SCP methods are still emerging. While some targeted antibody-based immunoassays have been applied to characterize proteins in single cells,^16,17^ these approaches are limited to a few dozen proteins per experiment and exhibit strong bias in quantification. Mass spectrometry (MS)-based proteomics can in principle overcome these limitations, but lacking the benefits of PCR, MS proteomic analysis at a single cell level is very challenging due to the extremely low amounts of proteins (ca. 0.2 ng in a mammalian cell), the high dynamic range of protein expression (7 orders of magnitude versus 3-4 orders for mRNAs),^18,19^ and the inevitable sample loss during protein extraction, digestion and chromatographic separation of the peptide digest.^20^ Consequently, despite the introduction of such ground-breaking SCP methods as SCoPE-MS^20^, SCoPE2^21,22^ and nanoPOTS^23^ that have been able to analyze between 500 and 2000 proteins in diverse cell lines, SCP has so far not been able to apply the techniques of chemical proteomics, such as FITExP. Here, we demonstrate such an ability, thus pioneering Single Cell Chemical Proteomics (SCCP).

Most SCP studies so far have considered two different types of cells (*e*.*g*., monocytes *vs* macrophage cells or Jurkat *vs* U-937 cells)^20,22^ with vastly different proteomes. Separation of these cells by SCP was relatively straightforward as it could be done using a few most abundant proteins. In contrast, in cells influenced by a drug the most significantly regulated proteins (drug targets) are seldom highly abundant, being frequently found in the abundance-sorted list below the 1000^th^ position. Therefore, the SCCP development required achieving following two intermediate objectives. First, average protein abundances in a homogeneous cell population measured by SCP must correlate with the abundances in bulk proteome analysis. This goal was achieved by starting from analyzing as bulk a relatively high number of cells and gradually reducing this number down to single cells, monitoring the correlation with the bulk analysis and systematically troubleshooting when this correlation broke down. A number of issues have been found and resolved related to protein extraction, digestion, labeling with isobaric regents, LC separation, MS acquisition and statistical analysis. At the end, satisfactory correlations between SCP and bulk proteomics results were consistently obtained. The second intermediate goal objective was to detect with SCP the known strong regulation of the drug targets, as in FITExP, with high statistical significance. This again required systematic studies and optimizations.

Here we present the SCCP workflow developed based on SCoPE-MS and applied to studying in a time-course manner the proteome effects of anticancer drugs MTX, camptothecin (CPT) and tomudex (TDX), also known as raltitrexed. These drugs were applied at LC_50_ concentration to A549 human lung adenocarcinoma cells, causing half of the cells to die within 48 h. Our workflow comprises the isolation of cells using fluorescence-activated cell sorting (FACS), minimal sample preparation including tryptic digestion, tandem mass tag (TMT) isobaric labeling for protein quantification, incorporation of a carrier proteome (CP) to boost the MS signal, chromatographic separation at a low flow rate, MS/MS data acquisitions and SCCP-optimized data processing (see **Figure 1**). The main goal of the study was to identify the time scale of the decision-making dying/surviving process, *i*.*e*., to reveal at what time the homogeneous cell population started to differentiate under the influence of a drug into cells committed to surviving or dying.

**Figure 1.**
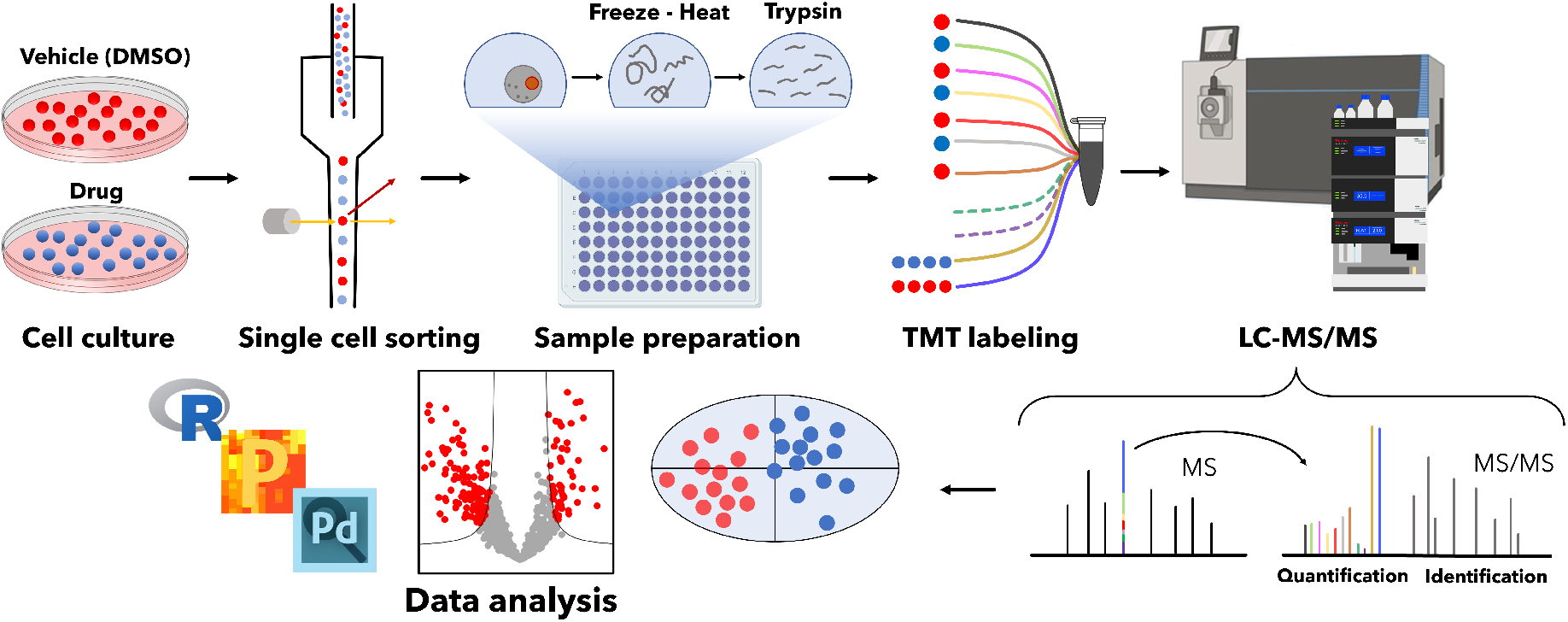
SCCP workflow. The workflow developed for SCCP included cell culturing and treatment with drugs, isolation of individual cells by FACS, protein extraction and digestion, TMT labeling of thus obtained tryptic peptides followed by multiplexing, LC-MS/MS and statistical data analysis. All steps are optimized for achieving the desired proteome depth and quantitative correlation with bulk analysis. In the figure, split carrier proteome occupies two channels (131N and 131C) in a TMT11plex set, with two other channels (130N and 130C) remaining empty (doted lines). Identification of peptides is achieved via matching masses of sequence-specific fragments, and quantification is performed by the abundances of the low-mass TMT reporter ions.

## 2. Results

### 2.1. Time-course SCCP analysis

The goal of the experiment was to determine the time point at which the attached cells make the decision to die, so that their proteome becomes altered to resemble that of the end-point detached (dying) cells rather than the end-point attached (surviving) cells. For that purpose, the cells were treated with MTX for 3, 6, 12, 24 and 48 h at LC_50_ concentration of 1.15 μM. The attached cells at each time point and the detached cells at 48 h were collected, and the FACS-isolated 96 cells of each type (**Figure S1**) were analyzed with SCCP using a single CP representing a mixture of the 48 h attached and 48 h detached cells. The bulk proteomes of 48 h detached and attached treated cells were analyzed separately. On average, over 1500 proteins and 10,000 peptides were identified and quantified in single cells at each incubation time. **Figure 2** shows how the attached treated and untreated cell populations, being almost indistinguishable on a PCA plot at 3 h treatment, become gradually separated with time, achieving nearly full separation at 12 h.

**Figure 2.**
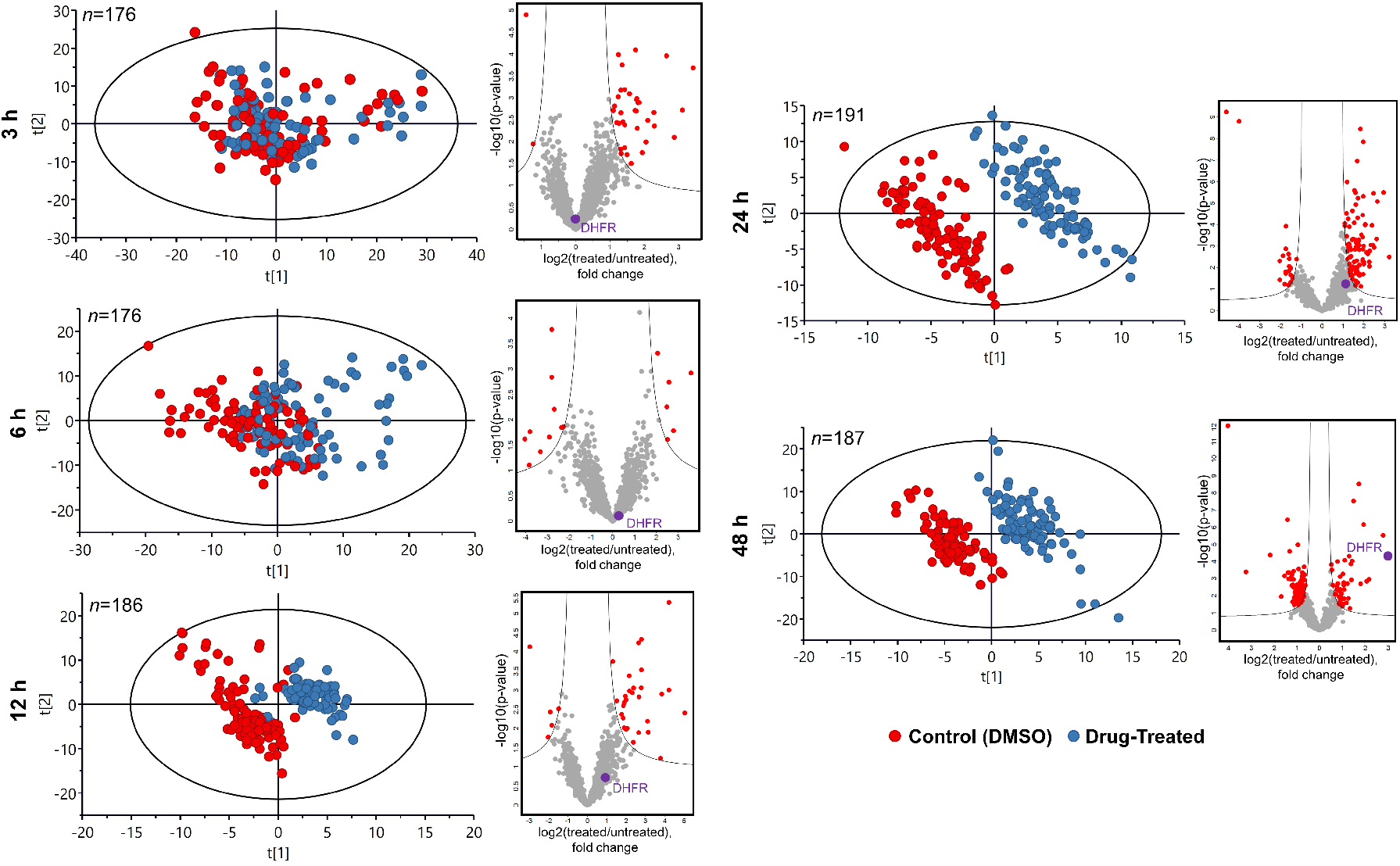
Time-course results upon treatment with MTX. PCA plots of SCCP data as a time course demonstrating the emergence of separation between the MTX-treated and untreated attached cells with incubation time, and the corresponding volcano plots of regulated proteins showing the emergence of DHFR among the top regulated proteins.

In order to identify at each timepoint the cell subgroup that was committed to death and the one committed to survival in the attached treated population, a hierarchical cluster analysis of protein abundances was performed (**Figure S2**). It was assumed that the two most abundant cell clusters represent the subgroups of the future surviving and dying cell. The hypothesis was that, being put on a PCA plot together with the 48 h attached and detached cells representing the two ultimate cell destinies, the two subgroups will reveal their identities by being closer to the respective destiny type. For time points earlier than the commitment event, cell clustering into the two subgroups will be random, and thus both subgroups would end up in the middle of the OPLS-DA plot close to each other.

Both these predictions were confirmed when the median abundances of all 1170 quantified proteins and 100 most abundant proteins in group 1 (G1) and group 2 (G2) separated by clustering analysis of attached cells were used for building an OPLS-DA model. The model also included the data on 48 h attached cells and 48 h detached cells, which represented the final destinations of the survival and dying subpopulations (**Figure 3A**). For 3 h and 6 h treatments, there was an overlap of the G1 and G2 clusters, with a separation between them in the direction of the destiny points at 12 h and longer treatment times. G1 was thus acquiring a proteome profile corresponding to the dying fate, while G2 represented the surviving subpopulation. As expected, the OPLS-DA separation between these two subpopulations grew with time. Similarly, the number of proteins with significantly changed abundances between the vehicle- and MTX-treated populations increased with time from 32 and 15 proteins at 3 and 6 h to 38, 121 and 134 proteins at 12, 24 and 48 h, respectively. Therefore, the A549 cell commitment to death occurs between 6 and 12 h past MTX treatment. This time scale is consistent with the earlier reports on dynamic proteomics measurements in cells treated with a drug at LC_50_ – in the first hours past treatment, the cells try to overcome the encountered difficulty, activating survival pathways, and only commit to death after such an attempt fails.^1^

**Figure 3.**
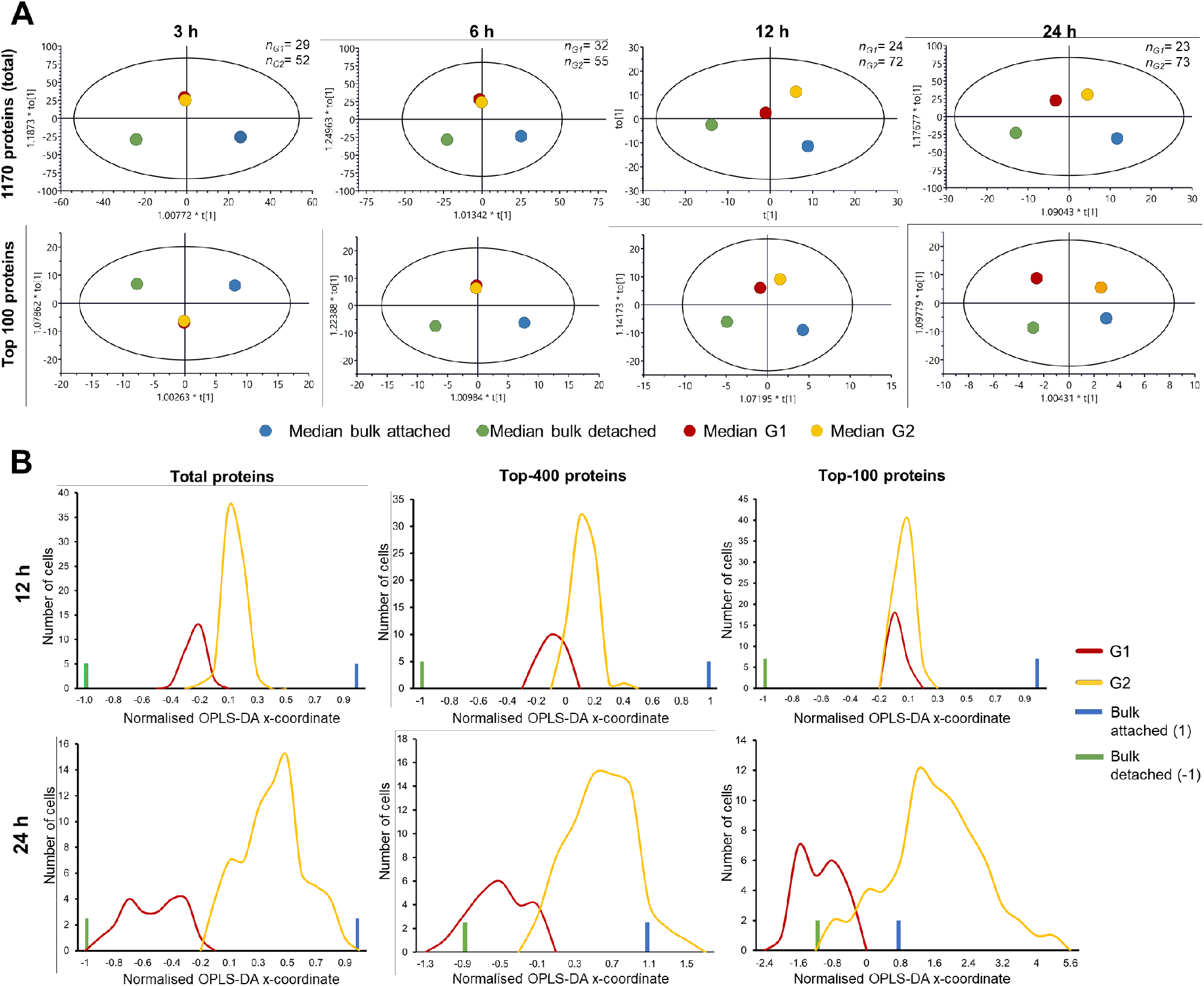
Statistical analysis of single cells treated with MTX. A) OPLS-DA analysis of SCCP data on median protein abundances in G1 and G2 cell groups from MTX treated cells at different time points together with bulk CP abundances for the total proteome (1170 proteins) and top 100 most abundant proteins. The numbers of cells belonging to G1 and G2 are given in the right top of each plot. B) Distribution of the main OPLS-DA coordinates of G1 and G2 groups of MTX-treated attached cells at 12 h and 24 h past treatment for total proteome, top 400 and top 100 proteins. The x-coordinates were normalized such that the coordinates of the attached and detached cell bulk-analyzed proteomes after 48 h treatment are +1 and -1, respectively.

Interestingly, at 12 h more separation was seen for the whole proteome, while at 24 h the 100 most abundant proteins showed bigger separation. This observation agreed well with the notion that the cell path to death starts with the inner mechanism altering lower-abundant mechanistic proteins first, followed by the altering household proteins that change the cell morphology. Consistent with this scenario, when the main OPLS-DA coordinates of individual cells were plotted on a scale normalized such that the attached cells treated for 48 h had x = 1 and the corresponding detached cells had x = -1, the obtained distributions of G1 and G2 cells were separated in 12 h for the full proteome, but less so for 400 most abundant proteins and not at all for top 100 proteins (**Figure 3B**). At the same time, for 24 h treatment the G1 and G2 proteomes gave broad distributions separated more for highly abundant proteins, suggesting that cell morphology alteration is well underway.

Pathway analysis of 179 proteins with significantly different abundances in G1 versus G2 at 12 h past MTX treatment revealed that they preferentially belong to metabolic, carbon metabolism, ribosome- and proteasome-related pathways (**Figure S3, Table S1)**. More specifically, the G1 subgroup is enriched in proteins involved in ribosome and proteasome-related pathways, meanwhile G2 subgroup is enriched in metabolic pathways.

### 2.2. SCCP with camptothecin (CPT) and tomudex/raltitrexed (TDX)

Similar results as with MTX were obtained with CPT (LC_50_ = 3 μM) and TDX (LC_50_ = 50 μM), with the targets TOP1 (downregulated) and TYMS (upregulated), respectively, emerging among the top proteins in the respective areas of the volcano plot (**Figure 4**). While these drugs have different MOAs and targets, the A549 cells have clearly formed two well separated clusters in PCA.

**Figure 4.**
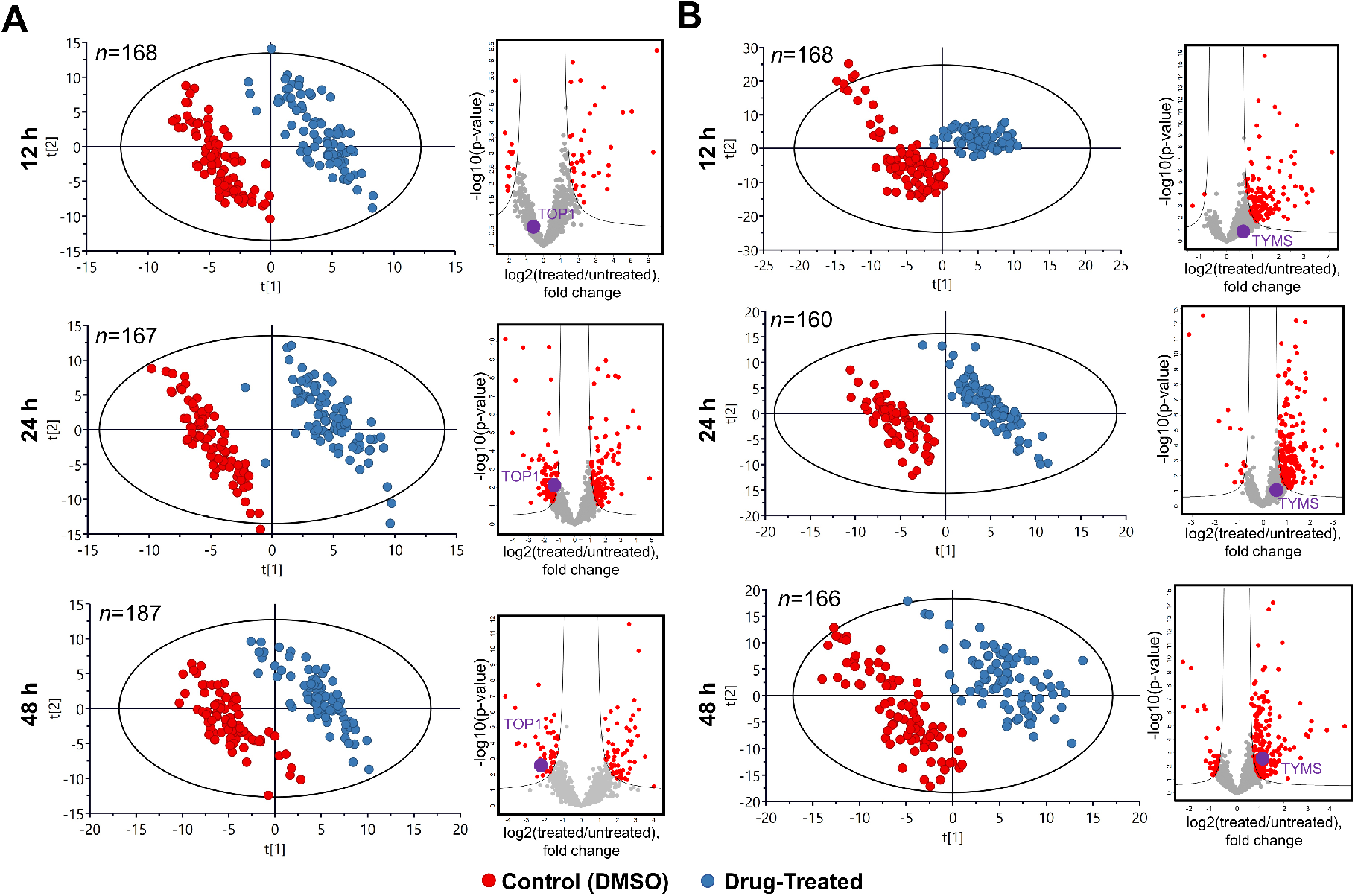
Time-course results upon treatment with CPT and TDX. PCA plots of SCCP data as a time course demonstrating the emergence separation between the untreated cells and the attached cells treated with A) CPT and B) TDX with incubation time, and the corresponding volcano plots of regulated proteins showing the emergence of the known drug target among the top regulated proteins.

### 2.3. Target percolation by OPLS-DA of MTX, CPT and TDX data

The ultimate goal of a Chemical proteomics experiment is drug target identification, which can be obtained by contrasting a specific treatment against all other treatments and controls. While designing ProTargetMiner^2^, we found that on average it takes 30-50 contrasting treatments to identify (“percolate”) the target uniquely among thousands of proteins in the proteome as the most specifically up- or down-regulated protein. Here, we merged the MTX, CPT and TDX SCCP data (treatment *vs*. untreated control) at 48 h of treatment and contrasted one drug against the other two (**Figure 5**). For MTX, the target DHFR was 4^th^ most specifically upregulated protein; for CPT, TOP1 was 15^th^ most specifically downregulated protein; and for TDX, TYMS was 10^th^ most specifically upregulated protein. These results demonstrate that SCCP has the potential for unique drug target identification, provided enough contrasting treatments are obtained.

**Figure 5.**
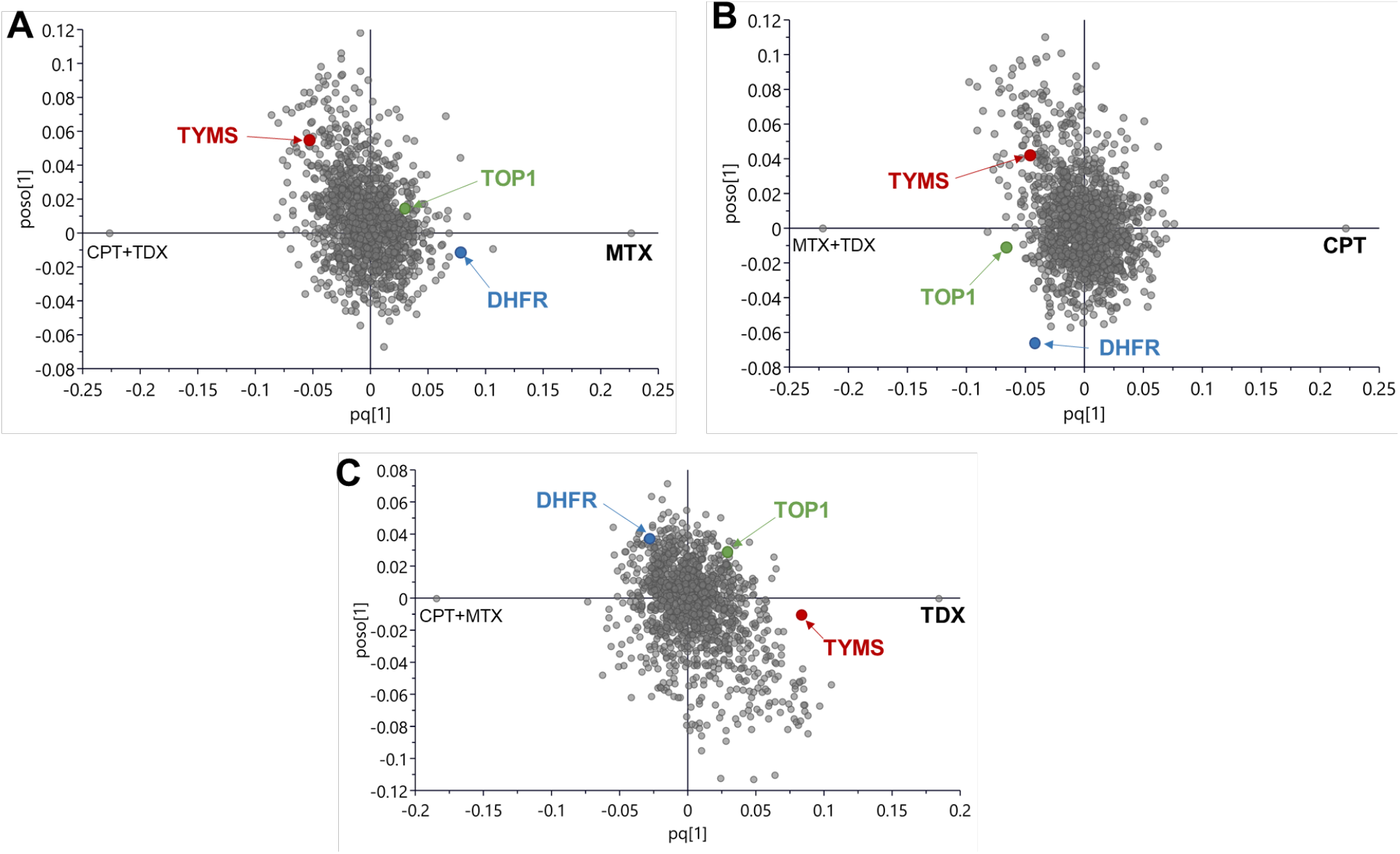
OPLS-DA analysis contrasting one drug (A) MTX, B) CPT and C) TDX) against the other two drugs, and the positions of the target proteins.

## 3. Discussion

Considering that cell to cell heterogeneity is a fundamental property of highly complex cellular systems^17^, the analysis of proteomes at a single cell level is essential for understanding the complex diseases, such as cancer, where diverse phenotypes contribute to the survival and progression,^24^ as well as for studying the mechanisms of cell resistance to anticancer treatment.

Here we demonstrated that SCP can be sufficiently quantitative for enabling Chemical proteomics approaches for drug target identification and monitoring. The ability to “percolate” by a contrasting OPLS-DA analysis, the probable drug target candidates has a paramount importance for the use of such a powerful drug target deconvolution method as ProTargetMiner.

SCCP time course analysis provided new biologically relevant information, confirming that cell commitment to death can now be studied at a proteome level for individual cells. Between 6 h and 12 h past treatment, a large group of attached drug-treated cells already committed to detach and form a floating dying population. Importantly, these changes were detected among the lower-abundant proteins, while highly abundant proteins remained at that point unaffected. It was even possible to determine the pathways and parts of cell machinery participating in the decision-making process.

## 4. Conclusions

After the quantitative aspect of single cell proteomics has been improved, chemical proteomics at the level of single cells became reality. The detailed profiling with SCCP of the heterogeneity of cancer cell response to drugs or treatments and the mechanistic analysis with cellular resolution of resistance to therapy is now possible. Moreover, the FITExP method of chemical proteomics is now applicable to single cells. As many novel analytical approaches, SCP is currently searching for the “killer application” that alone could justify this method and possess the capacity to dominate the applications. Exploring the heterogeneous response to drugs of cancer cells by SCCP might prove to be such a “killer application” for single cell proteomics.

The remaining challenges are however vast. For example, SCCP needs to provide deeper proteome analysis, targeting the benchmark of 5,000 proteins quantified with ≥2 peptides. A great achievement would be if complementary tools of chemical proteomics, such as proteome-wide integral solubility alteration (PISA) assay^25^ could be implemented for single cells. With PISA, one could monitor the protein target engagement of the drug molecule. This, however, requires significant efforts in improving the methods of handling and analyzing ultra-small protein amounts.

## 5. Methods

### 5.1. Cell culturing treatment

Human A549 lung adenocarcinoma cells obtained from ATCC (Manassas VA) were grown in Dulbecco’s Modified Eagle’s Medium (DMEM – Lonza, Wakersville MD) supplemented with 10% FBS superior (Biochrom, Berlin, Germany), 2 mM L-glutamine (Lonza) and 100 U/mL penicillin/streptomycin (Thermo, Waltham MA) at 37°C in 5% CO_2_.

The LC_50_ values for the drugs (MTX, CPT and TDX) were determined by CellTiter-Blue^®^ cell viability assay (Promega). Cells were seeded into 96 plates at a density of 3000 cells per well, and after 24 h of culture they were treated with serial concentrations of the respective drug: MTX (0 – 100 μM), CPT (0 – 100 μM) and TDX (0 – 100 μM). After 48 h, the media were discarded and replaced with 100 µL of fresh culture media. In each well, 20 µL of resazurin (CellTiter-Blue^®^ Cell Viability Assay kit – Promega) were added to perform the viability assay. After 4 h of incubation at 37°C, fluorescence of wells was measured in Infinite F200 Pro fluorometer (Tecan) by detecting the ratio between the excitation at 560 nm and emission at 590 nm. The LC_50_ values were determined from the dose-response curves by calculating the concentration causing the 50% fluorescence reduction compared with the untreated control.

Cells were then cultured and treated with MTX, CPT and TDX at LC_50_ concentrations in 75 cm^2^ flasks for 3, 6, 12, 24 and 48 h (for CPT and TDX only 12, 24 and 48 h treatment were performed). Control cells were treated with the vehicle (10 mM dimethyl sulfoxide - DMSO). After each incubation time point, the supernatant was collected and the attached cells were disconnected from the surface with TrypLE (Gibco) for 5 min, after which they were harvested by centrifugation at 1000 rpm for 3 min. Both types of cells (detached and adhered) were washed twice with cold 1X phosphate buffered saline (PBS).

### 5.2. Isolation of single cells by FACS

Collected attached cells were subjected to FACS analysis in FACSAria™ Fusion (BD Biosciences), in which cells were sorted based on the forward and side scatter (FSC/SSC) parameters only. Individual singlet cells were collected in a 96-well Lo-Bind plate (Eppendorf, Hamburg, Germany) containing 5 µL of 100 mM triethylammonium bicarbonate (TEAB) per well. A total of 96 single cells were sorted for each condition/drug (untreated and treated cells) using separate plates. In addition, a third plate was prepared, being dedicated only to CP. On that plate, 200 cells (100 treated plus 100 control) were collected per well in the first two rows. Altogether, 24 wells of CP cells were collected for each treatment and time point.

### 5.3. Protein extraction and digestion

Proteins from single cells and CPs were extracted in four freeze – thaw cycles. Plates were frozen for 2 min in liquid nitrogen and immediately heated at 37°C for 2 min. Proteins were denatured by heating the plates at 90°C for 5 min. The resulted protein solutions were centrifuged at 1000 rpm for 2 min to spin down all the volume present in the wells. Finally, 1 µL of 25 ng/µL sequencing grade trypsin (Promega, Madison WA) in 100 mM TEAB was dispensed using a MANTIS^®^ automatic dispenser (Formulatrix, Bedford MA). In the case of CP, digestion was achieved with 2 µL of trypsin solution added. Plates were incubated at 37°C overnight (16 h).

### 5.4. TMT labeling

TMT10plex™ and TMT11plex™ including channel 131C were used in this study. Unless specified, each TMT10plex™ set contained 4 control cells and 4 treated cells with tags interspaced, as well as a single channel with CP (200 cells in channel 131). TMT 130N was not used because of the cross contamination with the CP channel. Peptides were TMT-labeled by dispensing 1 µL of the respective TMT reagent dissolved in dry acetonitrile (ACN) at a concentration of 10 µg/µL using the MANTIS robot. Plates were incubated at room temperature (RT) for 2 h and then the reaction was quenched by adding 1 µL of 5% hydroxylamine (also with automatic liquid handler), following incubation at RT for 15 min. In some experiments, the CP was split in two channels, 131N and 131C, one composed of 100 control cells and the other one 100 treated cells. Channels 130N and 130C were left empty to prevent cross-contamination from CP channels. The labeled samples were pooled together using a 10-µL glass syringe (VWR, Japan), starting always with the CP samples in each TMT set in order to minimize sample loss during the pooling.^20^ Samples were pooled into MS-sample vials with glass insert (TPX snap ring vial from Genetec, Sweden) and dried in a speed vacuum concentrator (Concentrator Plus, Eppendorf). Dry peptides were resuspended in 7 µL of 2% ACN, 0.1% formic acid (FA) prior to LC-MS/MS analysis.

### 5.5. RPLC-MS/MS analysis

Peptide samples were separated on a Thermo Scientific™ Ultimate™ 3000 UHPLC (ThermoFisher Scientific) using a 10 min loading at 3 µL/min flow rate to a trap column (Acclaim™ PepMap™ 100, 2 cm × 75 µm, 3 µm, 100 Å - ThermoFisher Scientific). The separation was performed on an EASY-Spray™ C18 analytical column (25 cm × 75 µm, 1.9 µm, 300 Å – ES802A, ThermoFisher Scientific). A constant flow rate of 100 nL/min was applied during sample separation achieved in a linear gradient ramped from 5% B to 27% B over 120 min, with solvents A and B being 2% ACN in 0.1% FA and 98% ACN in 0.1% FA, respectively. LC-MS/MS data were acquired on an Orbitrap Fusion™ Lumos™ Tribrid™ mass spectrometer (ThermoFisher Scientific, San José CA), using nano-electrospray ionization in positive ion mode at a spray voltage of 1.9 kV. Data dependent acquisition (DDA) mode parameters were set as follows: isolation of top 20 precursors in full mass spectra at 120,000 mass resolution in the *m/z* range of 375 – 1500, maximum allowed injection time (IT) of 100 ms, dynamic exclusion of 10 ppm for 45 s, MS2 isolation width of 0.7 Th with higher-energy collision dissociation (HCD) of 35% at resolution of 50,000 and maximum IT of 150 ms in a single microscan. The mass spectrometry proteomics data are deposited to the ProteomeXchange Consortium via the PRIDE partner repository^26^ with the data set identifier PXD025481.

### 5.6. Data analysis

Raw data from LC-MS/MS were analyzed on Proteome Discoverer v2.4 (ThermoFisher Scientific), searching proteins against SwissProt human database (release July 30, 2019 with 20,373 entries) and known contaminants with Mascot Server v2.5.1 (MatrixScience Ltd, UK) allowing for up to two missed cleavages. Mass tolerance for precursor and fragment ions was 10 ppm and 0.05 Da, respectively. Oxidation of methionine, deamidation of asparagine and glutamine, as well as TMT adducts to lysine and N-termini were set as variable modifications. Percolator node^27^ in Proteome Discoverer was set to target false discovery rate (FDR) at 1% with validation based on *q*-value.

The TMT reporter ion abundances (RIA) at a peptide level were extracted from the search results. The subsequent analyses were performed in the RStudio (version 1.3.1073) programming language environment, the software for multivariate data analytics SIMCA (v. 15.0.2.5959, Sartorius) and Perseus software platform.^28^ Peptides from single cells with RIAs exceeding 10% of the abundance values for the respective carrier channel were filtered out, being considered a result of co-isolation or other interferences, resulting in about 30% of the peptides discarded for further analysis. After filtering, the remaining RIAs were arranged into a matrix of peptide IDs (rows) *vs*. single cells (columns). All RIAs were log2-transformed, and the data were normalized in columns by subtracting their median values computed, ignoring the missing values. Peptides quantified in less than ten cells were discarded (usually <0.05% peptides per dataset). Protein-level quantification was achieved by attributing each unique peptide to its respective top ranked protein within a protein group. As protein relative abundance, the median RIA value among the peptides belonging to that protein was taken.

The new relative abundance matrix (protein IDs *vs*. single cells) was again normalized by calculating the median value for each column (or single cell), and then, subtracting the median value calculated to each abundance on the respective column. Missing values in the resulting matrix were imputed based on the normal distribution of valid values (method available in Perseus software platform^29^), using a width of 0.3 standard deviations of the Gaussian distribution of the valid values and a downshift of 1.8 standard deviations. Finally, the batch effects across the TMT sets were corrected by applying an empirical Bayesian framework in the SVA package^30^.

The obtained matrix of relative protein abundances was used for statistical analysis. Principal component analysis (PCA) was performed to determine the separation degree between the control and drug-treated cells and to identify the outliers (single cells outside the limits of the PCA diagram with *p* < 0.05), which were removed from subsequent analysis. Resulting data were analyzed by orthogonal partial least squares discriminant analysis (OPLS-DA) and clustering analysis, and the fold changes were presented as volcano plots.

## 6. Acknowledgements

We would like to express our gratitude to Bogdan Budnik (Harvard Center for Mass Spectrometry, Harvard University, Cambridge MA) for discussion and advice on experimental design and suggesting the Mantis liquid handling robot, John Neveu (ESI Source Solutions, Woborn MA) and Erik Verschuuren (MS Wil B.V., Aarle-Rixtel, The Neatherlands) for ABIRD connection; Ujjwal Neogi and Flora Mikeloff (Karolinska Institutet) for valuable help with R algorithms, Indira Pla (Lund University) for useful advices in statistics and R algorithms, Willy Björklund, Erik Damgård and many others at Thermo Scientific for valuable technical support. The study was funded by the Swedish Research Council (grant 2018-06156, VR).

## 7. Authors contributions

RAZ conceptualized the study. AV and JER contributed to the conception and design of the study. AV and JER performed experiments and data analysis. RAZ wrote the manuscript and all authors contributed to the article and approved the submitted version.

