## Supplementary Figures for "Single Cell Chemical Proteomics (SCCP) Interrogates the Timing and Heterogeneity of Cancer Cell Commitment to Death"

**Figures S1 to S3**

Additionally, the processed mass spectrometric data used for analysis and interpretation is provided in form of tables compiled as compressed Excel file.

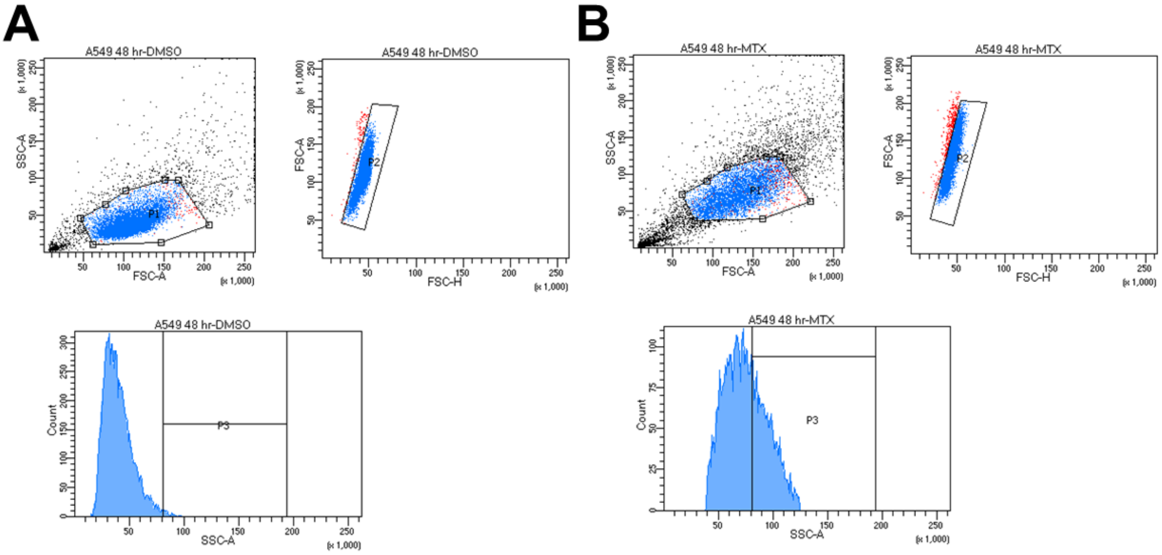


**Figure S1**. FACS gating conditions, isolating individual MTX DMSO treated (A) and treated (B) A549 cells after 48 h.

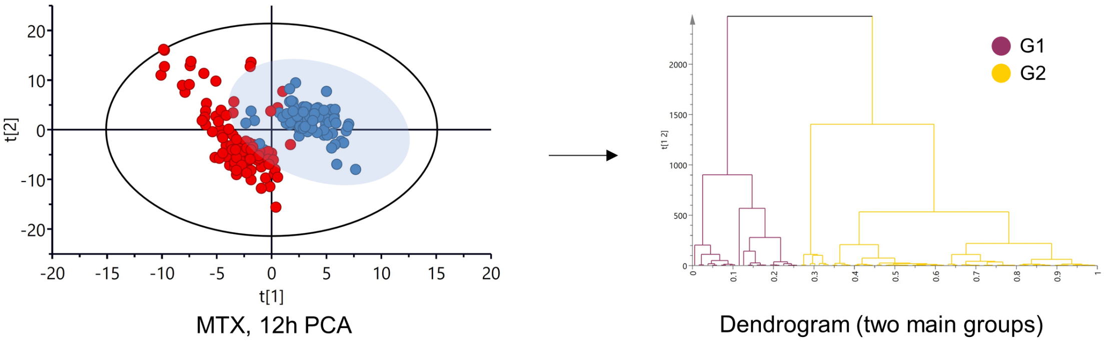


**Figure S2**. Clustering of the proteomes of attached treated single cells provides two groups of cells, G1 and G2.

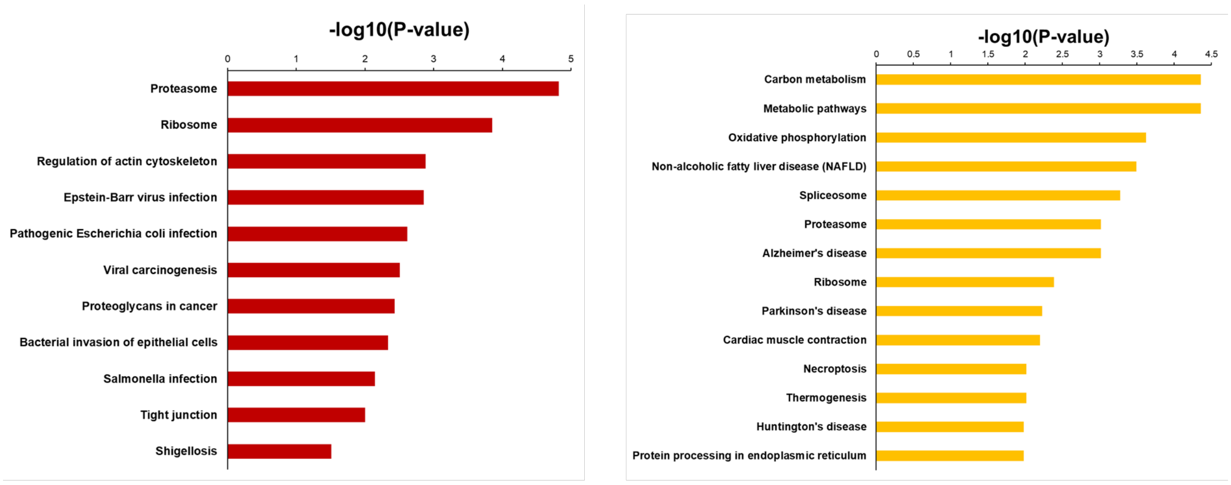


**Pathways enriched in G1 (101 proteins)** **Pathways enriched in G2 (78 proteins)**

**Figure S3**. Pathway analysis of 179 proteins with significantly different abundances in G1 versus G2 at 12 h past MTX treatment revealed that they preferentially belong to metabolic, carbon metabolism, ribosome- and proteasome-related pathways.
